# Prg4+ fibro-adipogenic progenitors in muscle are crucial for bone fracture repair

**DOI:** 10.1101/2025.05.14.654160

**Authors:** Qi He, Jiawei Lu, Qiushi Liang, Lutian Yao, Tingfang Sun, Huan Wang, Michael Duffy, Xi Jiang, Yuewei Lin, Ji-Hyung Lee, Jaimo Ahn, Nathaniel A Dyment, Foteini Mourkioti, Joel Boerckel, Ling Qin

**Author notes:** **Corresponding author:** Ling Qin **Email:**. **Author Contributions:** Qi He and Jiawei Lu contribute equally to this article. **Competing Interest Statement:** The authors declare no conflict of interest.

## Abstract

Clinically, compromised fracture healing often occurs at sites with less muscle coverage and muscle flaps can provide the necessary healing environment for appropriate healing in severe bone loss. However, the underlying mechanisms are largely unknown. Here, we established a mouse reporter model for studying muscle cell contribution to bone fracture repair. Analyzing skeletal muscle scRNA-seq datasets revealed that Prg4 marks a fibro-adipogenic progenitor (FAP) subpopulation. In mice, Prg4+ cells were specifically located in the skeletal muscle, but not at the periosteum or inside cortical bone. These cells expressed FAP markers, responded to muscle injury, and became periosteal cells under normal and muscle injury conditions. Fracture fragmented muscle fibers, rapidly expanded Prg4+ FAPs at the injury site and promoted their migration into the fracture site. Later, they gave rise to many chondrocytes, osteoblasts, and osteocytes in the outer periphery of callus next to muscle. In repaired bones, the descendants of Prg4+ FAPs were detected as mesenchymal progenitors in the periosteum and osteocytes at the prior fracture site. A second fracture activated those cells and stimulated them to become osteoblasts in the inner part of callus. Importantly, ablation of Prg4+ FAPs impaired fracture healing and functional repair. In an intramembranous bone injury model (drill-hole), Prg4+ FAPs became periosteal cells, but their contribution to bone defect repair was significantly less than in fractures. Taken together, we demonstrate the critical role of FAPs in endochondral bone repair and uncover a novel mechanism by which mesenchymal progenitors transform from muscle to cortical bone.

**Significance statement:** Fracture healing is often impaired in areas with less muscle coverage, but the mechanisms behind this are poorly understood. In this work, we uncovered a critical role of specific muscle cells, known as Prg4+ fibro-adipogenic progenitors (FAPs), in bone regeneration. Using mouse models, we show that these muscle-resident mesenchymal progenitors expand after bone injury, migrate to the fracture site, transform into various bone cells in the fracture callus, and eventually become bone surface progenitors after the bone is healed. This finding highlights the importance of muscle cells in endochondral fracture repair and reveals a novel crosstalk mechanism between muscle and bone tissues. Future studies targeting these muscle progenitor cells could develop new treatments for delayed and nonunion fractures.

## Introduction

Bone possesses remarkable regenerative healing capacity compared to most human tissues. After injury, bone and surrounding soft tissues undergo a well-coordinated process involving multiple cell types, cytokines, and growth factors—mimicking embryological processes—to restore bone to its pre-injured state (1). While this process is typically robust and efficient, about 5-10% of fractures can go awry and become delayed healing or even a nonunion (2). Factors that seem to modulate complication risk include age, diabetes, smoking, the anatomic site of the bone, the extent of soft tissue injury, infection, and fracture type. (3). Nonunion and subsequent reoperations place a considerable burden on patient’s functional capacity, quality of life, and healthcare resources. Currently, there are no FDA-approved pharmacological agents indicated to accelerate bone healing.

Every bone in the axial and appendicular skeleton is encased within a periosteum, a thin fibrous membrane pivotal for cortical bone expansion during growth and remodeling and for cortical bone regeneration after injury (4). It is well acknowledged that mesenchymal progenitors in the periosteum are the foremost important cell type for the functions of this thin layer (5). Injury promptly stimulates otherwise quiescent periosteal mesenchymal progenitors to proliferate and differentiate into chondrocytes and osteoblasts for callus formation. Removal of periosteum or damaging it by radiation eventually leads to fracture nonunion (6, 7). To date, a number of Cre drivers have been reported to label periosteal mesenchymal progenitors, including αSMA (8), Prx1 (9), Ctsk (10), Gli1 (7, 11), Pdgfrb (12), among others. Lineage tracing in reporter mice using these Cre drivers has helped characterize the crucial contribution of periosteal mesenchymal progenitors to callus formation and bone bridging following injury.

Meanwhile, clinical observation and practices have long suggested that nearby muscle also influences bone healing upon injury. For example, delayed healing and non-union rates are much higher in tibia fracture patients that had an associated compartment syndrome (13); compromised fracture healing is strongly associated with soft tissue injury, muscle viability, and coverage (14); muscle flaps in fracture patients with severe soft tissue injury assist in skeletal healing (15). An early animal study showed that muscle myogenic progenitors are present in the callus formed from open fracture with periosteum stripped, but not in the callus formed from closed fracture (16).

Using an elegant muscle graft transplant model, the Colnot group recently discovered that interstitial muscle fibro-adipogenic progenitors (FAPs) migrate to the fracture healing site and contribute to chondrocytes and osteoblasts in the callus (17, 18). However, it was unclear whether this observation is physiologically relevant as mice received additional surgery to implant exogenous muscle. Different from myogenic progenitors, FAPs are mesenchymal progenitors residing within muscle interstitium with the ability to differentiate into fibroblasts, adipocytes, and possibly osteoblasts, but not myoblasts (19–21). Research in the past decade has demonstrated that FAPs are crucial in supporting adult muscle homeostasis and regeneration following injury (22) and that they are the main source of ectopic bone formation in skeletal muscles (23). Despite their known importance in myogenesis and being the critical cell connecting muscle and bone, how FAPs interact with neighboring cells to shape biological processes remain largely unknown.

As mesenchymal progenitors, FAPs and periosteal progenitors share the same set of markers, such as Prxx1, Pdgfra, Sca1, and Gli1. This greatly hinders the researchers’ ability to directly investigate the role of FAPs in fracture under normal healing conditions without a transplant. Therefore, it became imperative to identify an FAP-specific marker that is not expressed in the periosteum. *Prg4* promoter-driven *CreER* was previously constructed to label the superficial layer of articular chondrocytes (24, 25). We recently constructed *Prg4-CreER Tomato* (*Prg4ER/Td*) mice to study the fate of synovial lining layer fibroblasts (26). Interestingly, we serendipitously observed Td labeling in muscle but not at periosteum. In the present study, we first demonstrated that Prg4+ muscle cells are FAPs and then used this reporter mouse line to unambiguously explore the fate and contribution of muscle-resident FAPs to cortical bone growth, maintenance, and repair. Two types of bone injury were utilized to study the extent of muscle cell contribution comprehensively: closed transverse fracture, which primarily heals by endochondral ossification, and cortical drill-hole defect, which mainly heals by intramembranous ossification. These studies not only demonstrate the critical role of Pdg4+ FAPs in skeletal regeneration but, importantly, they reveal a novel mechanism by which mesenchymal progenitors transform from muscle to cortical bone.

## Results

### Prg4 marks a FAP subpopulation within skeletal muscles but does not label bone cells

To locate Prg4-expressing cells and to trace their progenies in the musculoskeletal tissues, we crossed *Prg4-CreER* mice to td-Tomato (Td) reporter mice. These mice, *Prg4ER/Td*, at 2 months of age, were pulsed with daily Tamoxifen (Tam) injections on days 1-5 to induce gene recombination. Their hindlimbs were collected on day 8 for examination. Longitudinal sections showed that consistent with prior reports (24, 26) (Fig. 1A, Fig. S1A), Td+ cells resided at the superficial layer of articular cartilage, meniscus, and synovium (Fig. 1Ai). Notably, we observed some Td+ cells in the skeletal muscle surrounding the bone (Fig. 1Aii). However, we did not detect any Td+ cells in the periosteum and cortical bone (Fig. 1Aiii) nor inside the bone marrow (Fig. 1Aiv). To further evaluate this result, we performed RNA fluorescent in situ hybridization (FISH) on the hindlimbs of these mice. Similarly, we detected *Prg4* expression in muscle Td+ cells as well as Td+ cells at the surface layer of joint tissues (Fig. 1B, Fig. S1B, C).

**Figure 1.**
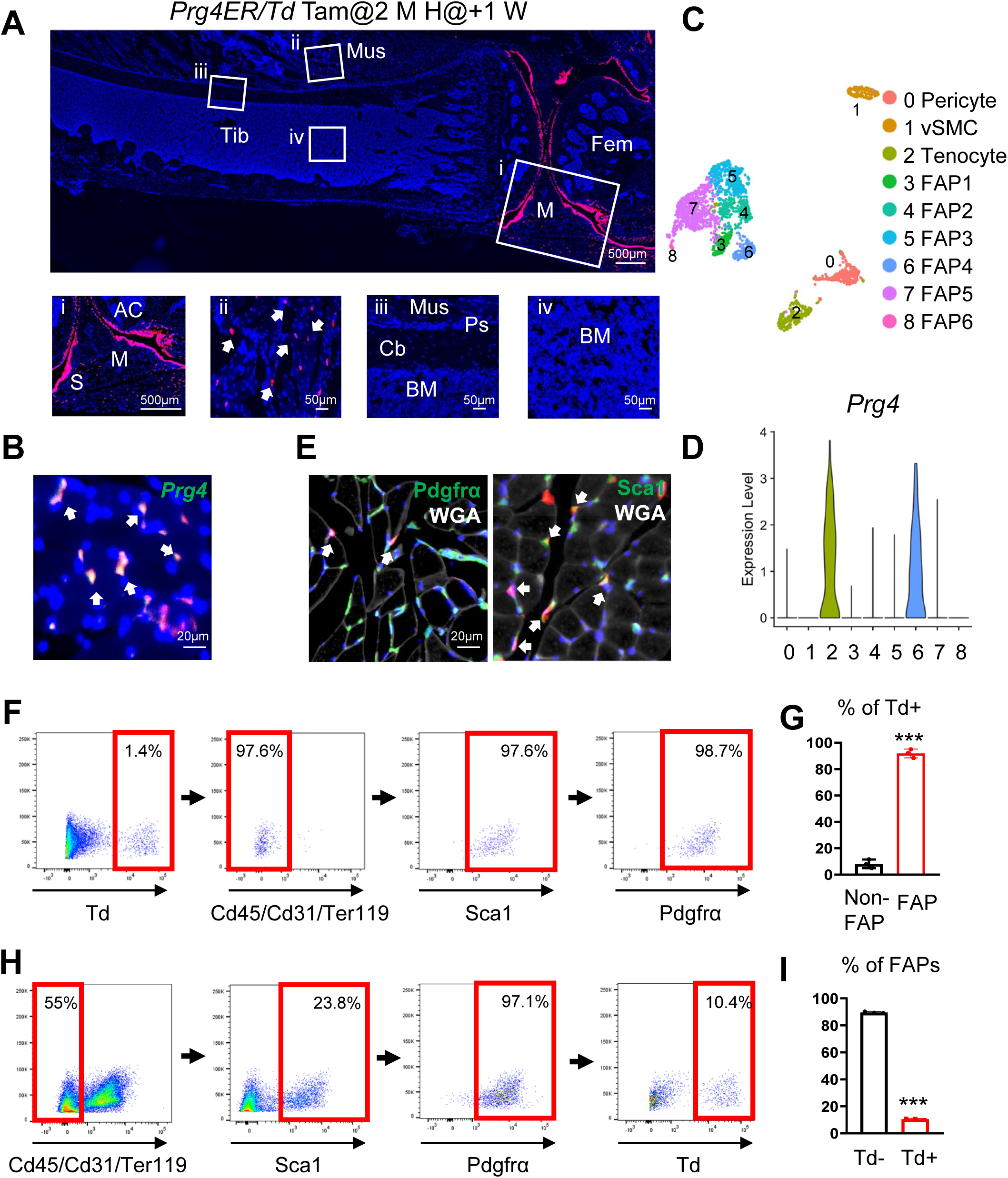
Prg4 expression labels a subset of FAPs in skeletal muscle. (A) A representative fluorescence image of a *Prg4ER/Td* hindlimb. Mice at 2 months of age received Tam injections for 5 days and their hindlimbs were harvested at day 7. Top panel: a low magnification image containing tibiae (Tib), femur (fem), and joint tissue. Scale bar: 500 μm. Bottom panel: squares in the top panel are magnified to show joint (i, scale bar: 500 μm), muscle (Mus, ii, scale bar: 50 μm), cortical bone (Cb, iii, scale bar: 50 μm), and bone marrow (BM, iv, scale bar: 50 μm). M: meniscus; S: synovium; AC: articular cartilage; Ps: periosteum. (B) Fluorescent image of TA muscle stained for *Prg4* mRNA by RNA FISH. Scale bar: 20 μm. (C) The Uniform Manifold Approximation and Projection (UMAP) plot of non-myogenic mesenchymal cells in mouse muscle. (D) Violin plot of Prg4 expression in muscle non-myogenic mesenchymal cell clusters. (E) Fluorescent images of TA muscle co-stained for Pdgfrα and WGA or Sca1 and WGA. Arrows point to Td+ Pdgfrα+ cells and Td+ Sca1+ cells. Scale bar: 20 μm. (F) Flow cytometer analysis of Td+ cells in muscle FAPs and non-FAPs. (G) Quantification of flow data. ***: p<0.001 vs Non-FAP, n=5 mice/time point. (H) Flow cytometer analysis of Td+ FAPs and Td-FAPs. (I) Quantification of flow data. ***: p<0.001 vs Td-, n=5 mice/time point.

To investigate which muscle cell types express Prg4, we analyzed publicly available single-cell and single-nuclei RNA-sequencing (scRNA-seq and snRNA-seq) datasets of skeletal muscle. In a mouse dataset (18), Prg4 was expressed in two clusters: FAP and Scx+ tendon cell (Fig. S2A). In a human dataset (27), Prg4 was specifically expressed in FAPs (Fig. S2B), consistent with another human scRNA-seq study identifying Prg4 as a marker of FAPs (28). Additionally, a recent mouse dataset focusing on non-myogenic mesenchymal cells further subdivided FAPs into 6 clusters (Fig. 1C) (29). Interestingly, Prg4 expression was restricted to a subcluster of FAPs (Clu1+ FAP4 cluster, Fig. 1D), which is associated with high mineralization ability, and a tenocyte cluster. Next, we stained *Prg4ER/Td* muscle with FAP markers Sca1 and Pdgfrα. Using fluorescently conjugated wheat germ agglutin (WGA) to visualize muscle fiber outlines, we found that Td+ cells are exclusively located in the interstitial area of myofibers, a location where FAPs reside (Fig. 1E). Among Td+ cells, 97.5+-0.2% are Sca1+ and 99.2+-0.7% are Pdgfrα+ (n=5 mice). Meanwhile, they were distinct from muscle stem cells (MuSCs), which are labeled by Myod or Pax7 (Fig. S3). To quantify Prg4+ cells in FAPs, we digested anterior and posterior tibial muscle for flow cytometry analysis. Td+ cells constituted 1.4% of total cells, and they were lineage negative (Lin-: CD31-CD45-Ter119-) and Sca1+Pdgfrα+ (Fig. 1F, G). Among Lin-Sca1+Pdgfrα+ FAPs, 10.4% were Td+ (Fig. 1H, I). Colony-forming unit-fibroblast (CFU-F) assay of digested muscle cells showed that approximately 23.6% of CFU-Fs are Td+ (Fig. S4). Collectively, these data indicate that Prg4+ cells are a subpopulation of FAPs residing within skeletal muscle.

### Muscle FAPs become periosteal cells over time under physiological condition but not vice versa

To trace the cell fate of Prg4+ FAPs in growing mice, we injected Tam into 2-week-old *Prg4ER/Td* mice and harvested hindlimbs at 1, 4, and 12 weeks later for analysis (Fig. 2A, B). During this period, the total number of Td+ cells increased over time, but their density was not altered. While we did not detect Td+ cells inside the periosteum at 1 and 4 weeks post induction, some Td+ cells appeared in the periosteum at 12 weeks. We next injected Tam into 2-month-old *Prg4ER/Td* mice (Fig. 2C, D). Throughout the tracing period, muscle volume did not change significantly, and Td+ cells remained unaltered. Similarly, we observed Td+ cells in the periosteum at 12 weeks. Periosteum is composed of inner cambium layer and outer fibrous layer. Osterix specifically labels osteogenic cells in the cambium layer (30). Interestingly, Td+ cells were Osterix-but in close proximity to Osterix+ periosteum cells (Fig. S5), suggesting that they reside in the fibrous layer. These data provide the first evidence that Prg4+ FAPs have the ability to become periosteal bone cells.

**Figure 2.**
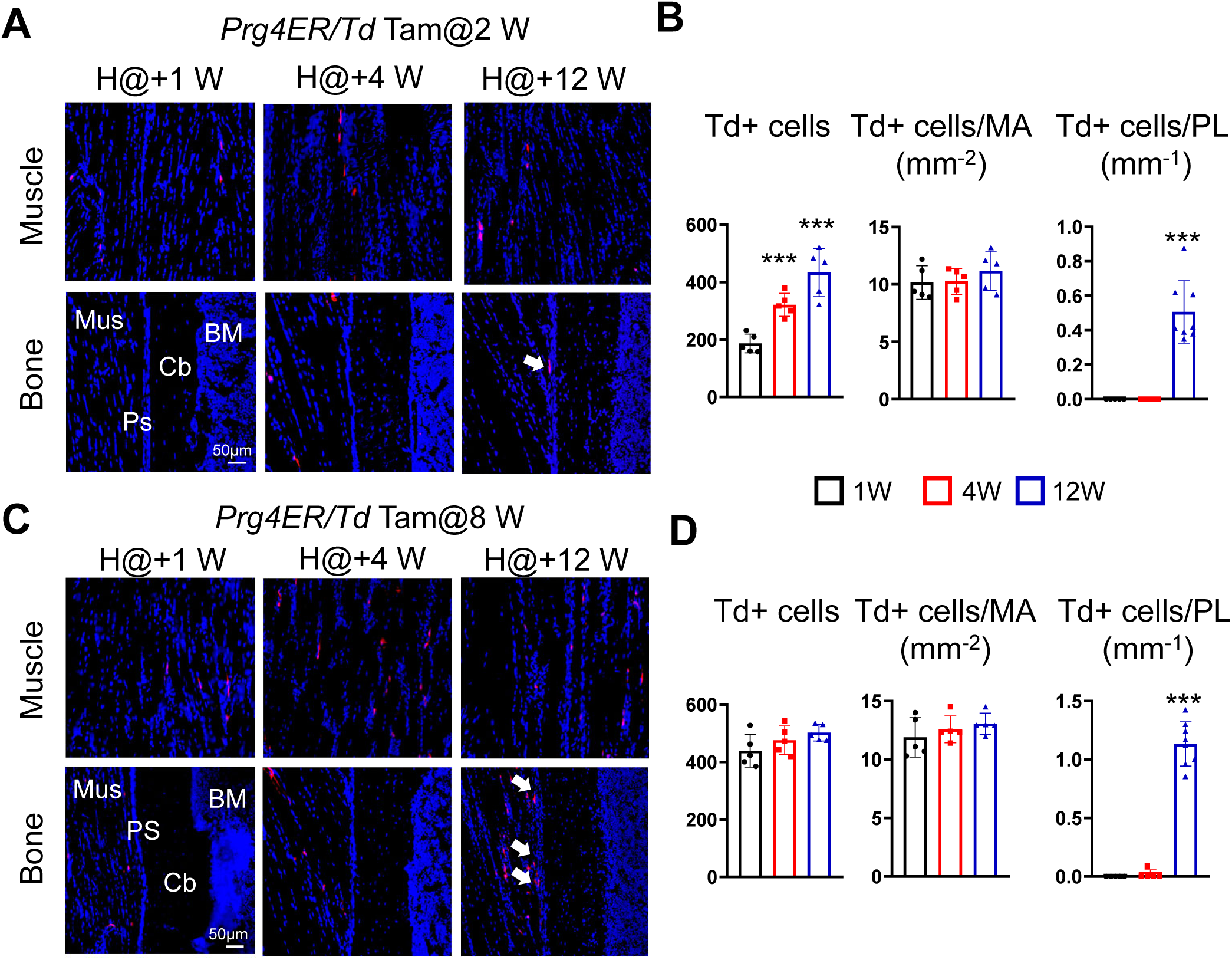
Prg4+ FAPs give rise to periosteal cells during muscle growth and maintenance. (A) Fluorescence images of *Prg4ER/Td* tibiae and surrounding muscle harvested at 1, 4, and 12 weeks after Tam injections. Mice received Tam at 2 weeks of age. Mus: muscle; Cb: cortical bone; Ps: periosteum. Arrows point to Td+ cells in the periosteum. Scale bar: 50 μm. (B) Total Td+ cells in muscle, the percentage of Td+ cells in muscle, and the number of Td+ cells in the periosteum were counted. MA: muscle area. PL: periosteum length. ***: p<0.001 vs1 week, n=5 mice/time point. (C) Fluorescence images of *Prg4ER/Td* tibiae and surrounding muscle harvested at 1, 4, and 12 weeks after Tam injections. Mice received Tam at 8 weeks of age. Arrows point to Td+ cells in the periosteum. Scale bar: 50 μm. (D) Total Td+ cells in muscle, the percentage of Td+ cells in muscle, and the number of Td+ cells in the periosteum were counted. ***: p<0.001 vs 1 week, n=5 mice/time point.

To assess whether bone cells could become muscle cells, we examined hindlimbs of 2-month-old *Col2-Cre Td* (*Col2/Td*) mice. Our previous reports demonstrated that Td labels the entire lineage of mesenchymal cells in periosteum and bone marrow, including mesenchymal progenitors and terminally differentiated cells (31, 32). Examining tibiae along with muscle confirmed that Td labels periosteum and osteocytes in the cortical bone (Fig. S6). However, we did not detect any Td+ cells in nearby muscle tissue, suggesting that bone cells do not become muscle cells during normal growth.

### Prg4+ FAPs expand rapidly after acute muscle injury and give rise to periosteal cells

FAPs are well known for their responses to acute muscle injury and their critical role in muscle repair (33). To examine whether Prg4+ FAPs have similar properties, we applied two types of muscle injury on 2-month-old *Prg4ER/Td* mice at 2 weeks after receiving Tam injections. In the first model, mice were intramuscularly injected with 50% glycerol. Six days later, H&E staining revealed modest muscle degeneration (Fig. 3A). Fluorescent images showed a great expansion of Td+ cells in the muscle (Fig. 3B), which was further confirmed by a 5.3-fold increase in the flow analysis (Fig. 3C). Glycerol injection induces muscle adiposity and FAPs are the cell source for these adipocytes (34). We observed that some newly formed adipocytes in muscle are Td+ (Fig. 3D), suggesting that Prg4+ FAPs differentiate into adipocytes after injury. Interestingly, we noticed many Td+ cells in the fibrous layer next to Osterix+ cells in the cambium layer after glycerol injection (Fig. 3E), further supporting that Prg4+ FAPs can become periosteal bone cells, in this case, in response to injury.

**Figure 3.**
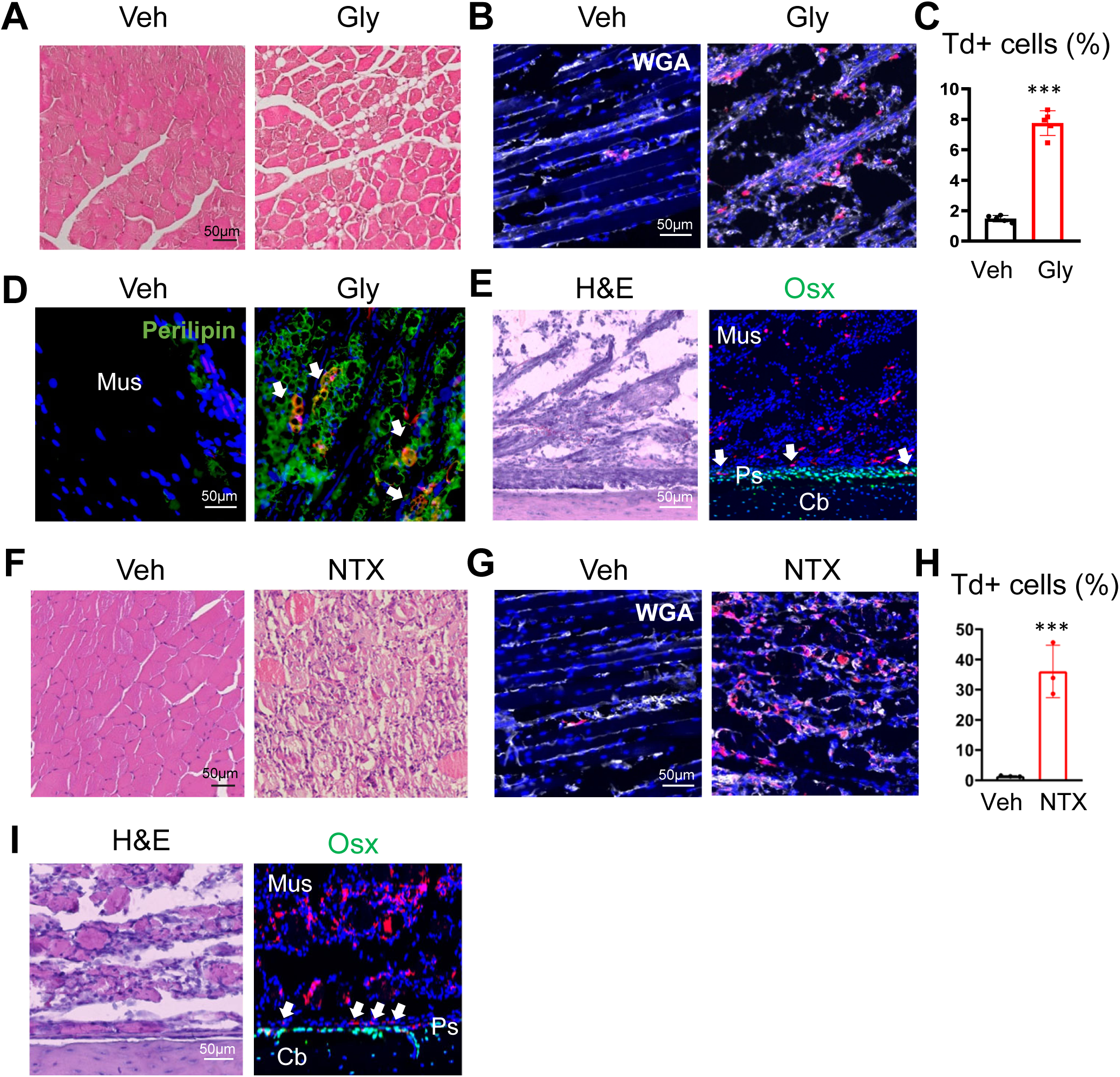
Prg4+ FAPs respond drastically toward muscle injuries. (A) Representative H&E images of TA muscle from *Prg4ER/Td* mice at 6 days after receiving a vehicle (veh) or a glycerol (Gly) injection. Mice at 2 months of age received Tam injection for 5 days at 2 weeks before muscle injury. Scale bar: 50 μm. (B) Fluorescence images of *Prg4ER/Td* TA muscle. Scale bar: 50 μm. (C) The percentage of Td+ cells among Lin-cells was quantified. ***: p<0.001 vs Veh, n=5 mice/group. (D) Fluorescence images of *Prg4ER/Td* TA muscle stained by Perilipin to show adipocytes (arrows). Mus: muscle. Scale bar: 50 μm. (E) H&E and fluorescence images with Osterix straining of adjacent tibial sections from *Prg4ER/Td* mice after injury. Arrows point to Td+ cells in the fibrous layer. Mus: muscle; Cb: cortical bone; Ps: periosteum. Scale bar: 50 μm. (F) Representative H&E images of TA muscle from *Prg4ER/Td* mice at 3 days after receiving a vehicle (veh) or a notexin (NTX) injection. Mice at 2 months of age received Tam injection for 5 days at 2 weeks before muscle injury. Scale bar: 50 μm. (G) Fluorescence images of *Prg4ER/Td* TA muscle. Scale bar: 50 μm. (H) The percentage of Td+ cells among Lin-cells was quantified. ***: p<0.001 vs Veh, n=3 mice/group. (I) H&E and fluorescence images with Osterix straining of adjacent tibial sections from *Prg4ER/Td* mice after injury. Arrows point to Td+ cells in the fibrous layer. Mus: muscle; Cb: cortical bone; Ps: periosteum. Scale bar: 50 μm. A, D, F: samples were cut in a cross-sectional plane. B, E, G, I: samples were cut in a longitudinal plane.

In the second model, mice were intramuscularly injected with notexin (NTX), a myotoxin widely used for studying muscle injury and repair. NTX caused severe muscle degeneration at day 3 (Fig. 3F) and a great expansion of Td+ cells in muscle (Fig. 3G). Flow analysis validated a 25.7-fold increase of Td+ cells among Lin-muscle cells (Fig. 3H). Similarly, we also noticed the appearance of Td+Osterix-periosteal cells after NTX injury (Fig. 3I). Overall, these results suggest that Prg4+ FAPs undergo dynamic changes upon acute muscle injury.

### Muscle-resident, but not tendon-resident, Prg4+ cells contribute to fracture callus formation

In a closed, transverse fracture model, bone injury is achieved by weight drop, which inevitably damages muscle surrounding bone, resulting in many small muscle fragments at the injury site (Fig. 4A). Since Prg4+ FAPs expand after muscle injury, we studied their responses toward fracture injury and traced their fate in fracture healing. For this experiment, we injected Tam into 2-month-old *Prg4ER/Td* mice and performed a close fracture on tibiae 2 weeks later. Fractured tibiae were collected on days 3, 7, 14, 28, and 56 to represent distinct stages of fracture healing (Fig. S7). On day 3 when the periosteum thickened, we observed abundant Td+ cells in the fracture hematoma, as well as a drastic increase of Td+ cells in the muscle around the injury site (Fig. 4Bi, ii, Fig. S8), indicating that Prg4+ FAPs expand in respond to bone injury and their descendants quickly migrate to the fracture gap. Cd45 staining showed that those cells are not hematopoietic cells. In the meantime, the periosteum close to the fracture gap thickened but did not contain any Td+ cells (Fig. 4Biii), suggesting that periosteal mesenchymal progenitors are the major contributor to the expansion of periosteum immediately post fracture. Periosteum distal to the injury site remained Td-(Fig. 4Biv).

**Figure 4.**
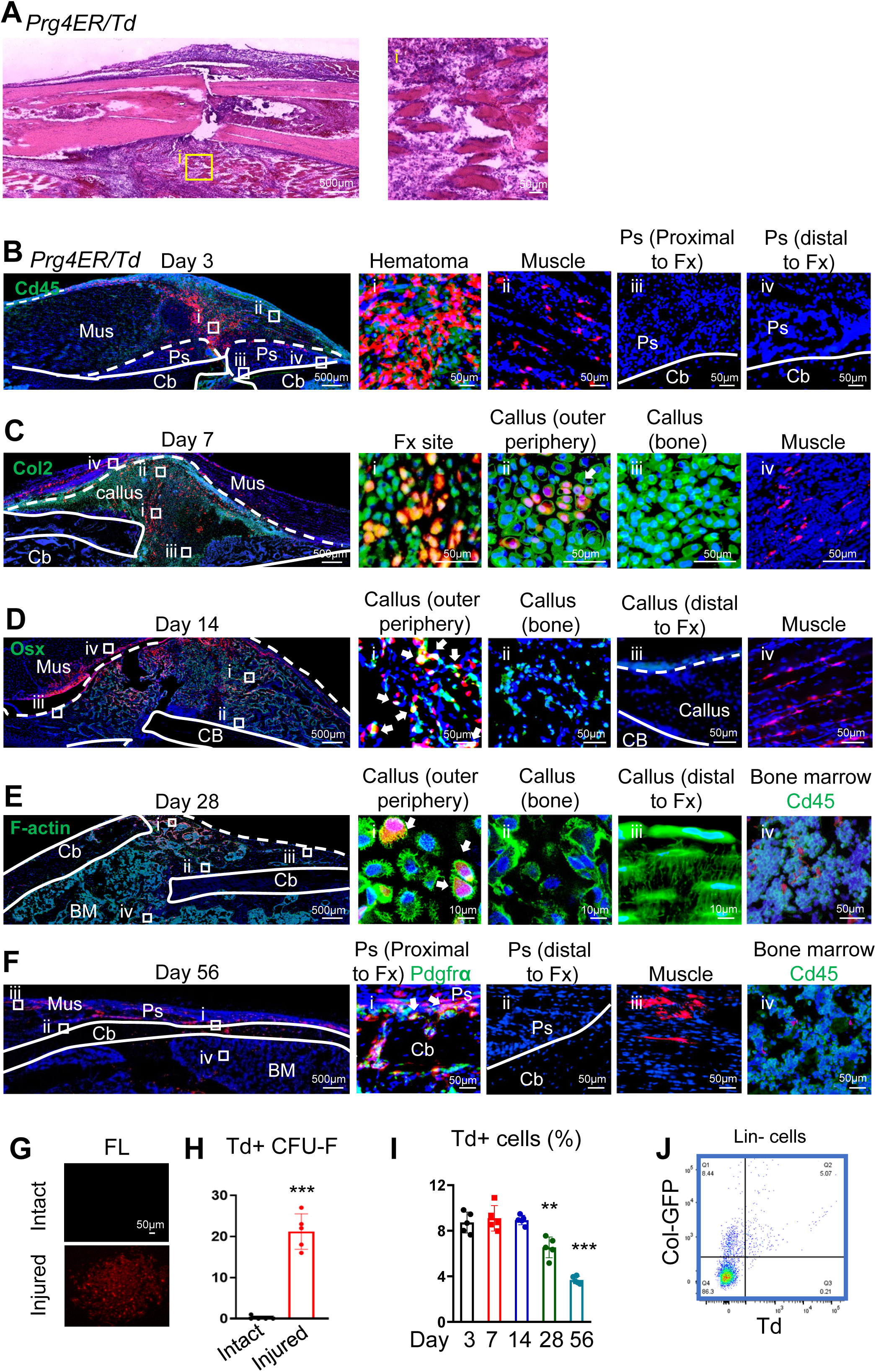
Prg4+ FAPs contribute to fracture healing. (A) H&E staining of fractured bone and surrounding muscle at 3 days post fracture. Scale bar: 500 μm. (i, scale bar: 50 μm), (B-F) Representative fluorescence images of tibial fracture samples from *Prg4ER/Td* mice harvested at days 3 (B), 7 (C), 14 (D), 28 (E), 56 (F) post surgery, co-stained with Cd45 (day 3), Col2 (day 7), Osterix (Osx, day 14), and phalloidin (F-action, day 28). Mice at 2 months of age received Tam injections for 5 days and fracture surgery at right tibiae 2 weeks later. The left panels are low magnification images (scale bar: 500 μm), and the right panels are high magnification images of squares in the left panels (scale bars are listed in the images). In Di, Ei, Fi, arrows point to Td+Osterix+ osteoblasts, Td+ osteocytes, and Td+Pdgfra+ periosteal cells, respectively. Mus: muscle; Cb: cortical bone; Ps: periosteum; BM: bone marrow. (G) Fluorescence images of CFU-F colonies from the peristeum of intact bone and bone at day 56 post fracture. (H) CFU-F numbers were counted from 1 million digested periosteal cells. n=5 mice/group. (I) The percentage of Td+ cells among Lin-cells from fracture callus was measured by flow cytometry over time. **: p<0.01, ***: p<0.001 vs day 3, n=5 mice/group. (J) Flow analysis of Td+ and GFP+ cells from day 28 fracture callus in *Prg4ER/Td*/*Col1GFP* mice. n=5 mice/group.

On day 7, when a soft callus was formed, many type II collagen (Col2+) chondrocytes close to the fracture end or in the outer periphery of the callus were Td+ (Fig. 4Ci, ii), indicating that muscle FAPs differentiate into chondrocytes and become a part of fracture callus. In contrast, cells in the callus close to the cortical bone were Td-(Fig. 4Ciii), suggesting that they likely come from periosteal cells. Nearby muscle still contained an increased amount of Td+ FAPs (Fig. 4Civ).

On day 14, when a hard callus was formed, a lot of osteoblasts, labeled by Osterix staining, in the outer periphery of the callus were Td+ (Fig. 4Di). However, those in the callus close to bone or distal to the fracture site, were Td-(Fig. 4Dii, iii). Nearby muscle still has expanded Td+ cells (Fig. 4Div).

On day 28, by using F-actin staining to identify osteocytes in the callus, we found that Prg4-lineage cells become osteocytes at similar locations as osteoblasts (Fig. 4Ei-iii). Interestingly, we also observed some Td+Cd45-cells inside the bone marrow (Fig. 4Eiv). This is in sharp contrast to the intact bone with no Td+ cells in the bone marrow.

On day 56, when the bone was almost healed, we observed that the regenerated periosteum and cortical bone at the prior fracture site contained Td+ cells (Fig. 4F). Most of them were Pdgfrα+ (Fig. 4Fi), a marker for periosteal mesenchymal progenitors. However, periosteal cells distal to the injury site were Td-(Fig. 4Fii). Meanwhile, we still observed increased Td+ cells in nearby muscle (Fig. 4Fiii). Similar to day 28, bone marrow within the newly formed cortical bone contained Td+ cells (Fig. 4Fiv). CFU-F assay of periosteal cells revealed Td+ colony formation from fractured bones but not from intact bones (Fig. 4G, H), further demonstrating that muscle-resident Prg4+ FAPs become periosteal mesenchymal progenitors after injury.

To quantify the contribution of Prg4-lineage cells to fracture callus, we examined cells dissociated from callus at various healing stages by flow cytometry. Td+ cells comprised 8.7%, 9.1%, 8.9%, 6.5%, and 3.6% of Lin-callus cells at day 3, 7, 14, 28, and 56, respectively, post fracture (Fig. 4I). Note that they are underestimated because we harvested fracture callus together with bone marrow inside cortical bones. To quantify the contribution of Prg4-lineage cells to bone-forming cells in the callus, we incorporated *Col13.6-GFP*, a marker for osteoblasts and osteocytes, into *Prg4ER/Td* mice and applied similar Tam administration and fracture on these mice. Flow analysis of day 28 fracture showed that 34.3+-4.3% of GFP+ cells are Td+, and 90.7+-10.9% of Td+ cells are GFP+ (Fig. 4J).

Since Prg4 labels not only muscle FAPs but also tendon cells (Fig. 1C, D), we next investigated whether tendon cells participate in fracture healing. To do so, we examined fracture healing in two mouse models containing reporters driven by tendon-specific *Scx* promoter. Analyzing hindlimb samples from *Scx-GFP* mice revealed that, as expected, tendons appear within the muscle but not at the long bone midshaft, the fracture site (Fig. S9A). Using adjacent sections for brightfield and fluorescence imaging, we observed that, unlike muscle being damaged into small fragments, tendons remain largely intact after fracture (Fig. S9B, C). We also generated *Scx-CreER Td* (*ScxER/Td*) mice for lineage tracing. After tam injections, tendon at the muscle/bone junction was labeled by Td (Fig. S10A). Two weeks later, these mice received a tibial fracture. Very few Td+ cells were detected in the hematoma on day 3 (Fig. S10B), and almost no Td+ cells appeared in the fracture callus on day 14 (Fig. S10C), suggesting that tendon-derived cells do not contribute to callus formation. After excluding the participation of tendon cells, our results unambiguously demonstrated a significant contribution of muscle-resident Prg4+ FAPs to fracture callus and revealed their multi-lineage differentiation abilities into chondrocytes and osteoblasts.

### Prg4+ FAP-derived periosteal cells respond to the next fracture injury

To further explore whether the descendants of Prg4+ FAPs in periosteum are mesenchymal progenitors, we subjected *Prg4ER/Td/Col1GFP* mice to fracture at 2 weeks after Tam injections, and another fracture at the same site at day 56 post the first surgery. On day 3 post the second fracture, we observed many Td+ cells in the fracture hematoma and in the thickened periosteum (Fig. 5A). Similarly, on day 14 post the second fracture, Td+ osteoblasts were present throughout the callus, not only in the area close to muscle but also in the area close to the bone (Fig. 5B). These results showed that Td+ cells in the periosteum, which are the descendants of Prg4+ FAPs after the first injury, can be activated for expansion upon the next injury.

**Figure 5.**
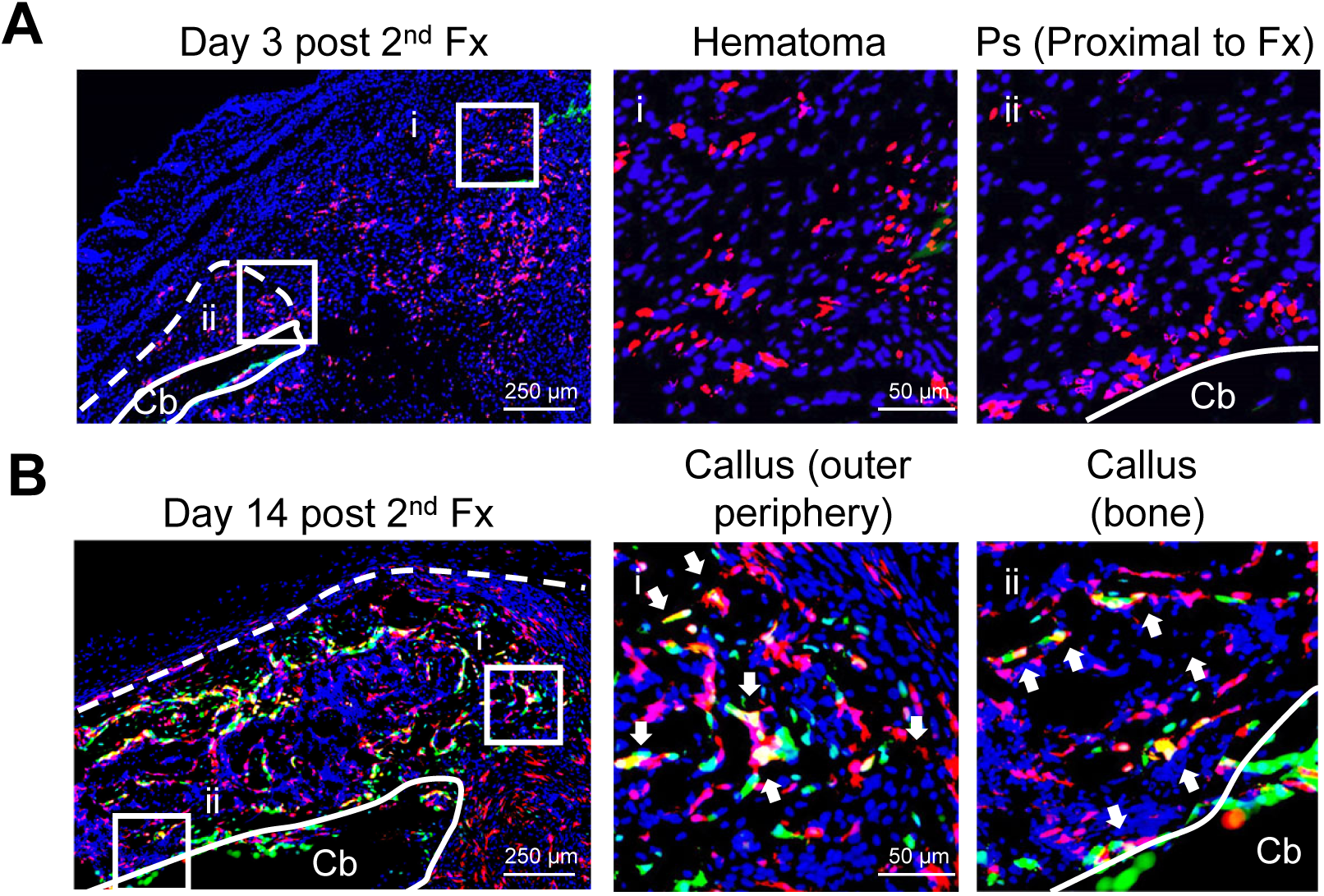
Prg4+ FAP-derived periosteal mesenchymal progenitors respond to a second fracture. Representative fluorescence images of tibial fracture samples from *Prg4ER/Td/Col1GFP* mice on day 3 (A) and day 14 (B) after the second fracture. Mice at 2 months of age received Tam injections for 5 days and the first fracture 2 weeks later. At day 56 post the first surgery, mice received the second fracture at the prior fracture site. The left panels are low magnification images (scale bar: 250 μm). Squares from those images are magnified at the right panels (scale bar: 50 μm).

### Prg4+ FAP ablation impairs fracture healing

To access the impact of Prg4+ FAPs on bone regeneration, we depleted these cells using a mouse model overexpressing DTA (diphtheria toxin A-subunit). *Prg4-CreER Td DTA* (*Prg4ER/DTA*) and *WT* control (*Prg4ER/Td*) mice were subjected to Tam inductions for 5 days at 2 months of age. Three days after the last injection, histology revealed a great reduction of Td+ cells in the muscle and joint of DTA mice (Fig. 6A, S11) and flow cytometry revealed a 62.8% decrease of Td+ cells in muscle Lin-cells (Fig. 6B, C), confirming the success of cell ablation. These mice were subjected to fracture at 2 weeks after Tam injections. MicroCT analysis revealed a significant reduction of bone volume (BV) and bone volume fraction (BV/TV) in the callus at day 14 and day 21 post fracture (Fig. 6D-F). At day 56, while fractured bones were all repaired in *WT* mice, only 40% were bridged in *Prg4ER/DTA* mice, leading to a decreased healing score (Fig. 6D, G). Moreover, bone mechanical strength, characterized by maximum torque, torsional stiffness, and torsional energy to failure, was also significantly reduced in *Prg4ER/DTA* mice (Fig. 6H). Together, these findings indicate that Prg4+ FAPs are essential for fracture callus formation and functional repair.

**Figure 6.**
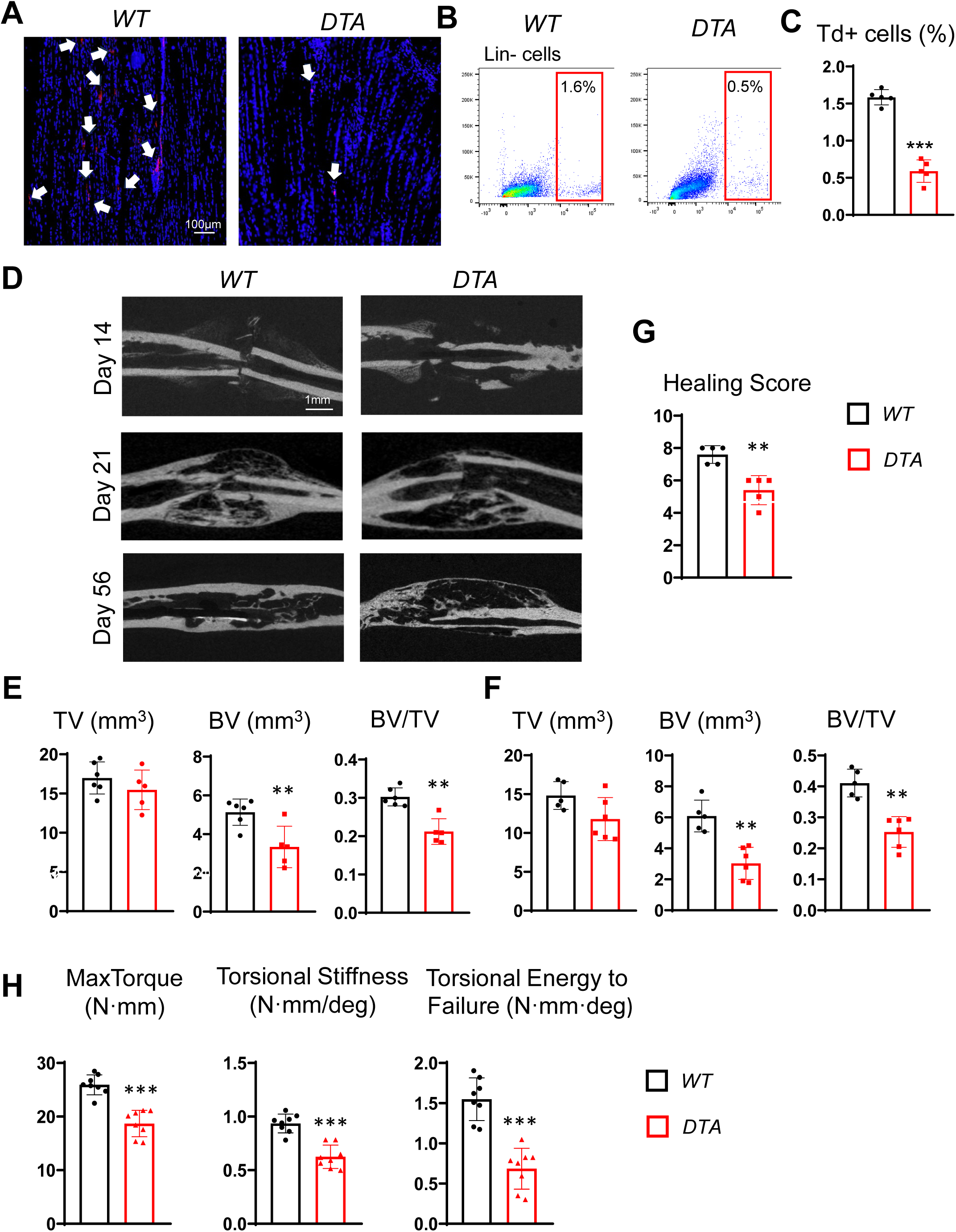
Ablation of Prg4+ FAPs cause delayed fracture healing. (A) Fluorescence image of *Prg4ER/Td* (*WT*) and *Prg4ER/Td/DTA* (*DTA*) muscle. Arrows point to Td+ cells in muscle. Scale bar: 100 μm. (B) Flow analysis of Td+ cells among Lin-cells in muscle. (C) Quantification of the percentage of Td+ cells in Lin-cells. n =5 mice/group. (D) Representative 2D micro-CT scan image of *WT* and *DTA* tibiae on days 14, 21, and 56 post fracture surgery. Scale bar: 1mm. (E, F) MicroCT measurement of fracture callus structural parameters on day 14 (E) and day 21 (F) post fracture. TV: tissue volume; BV: bone volume; BV/TV: bone volume fraction. n =5 mice/group. (G) Fracture healing scores were quantified at day 56 post fracture. n =5 mice/group. (H) Mechanical testing was performed on bones at 6 weeks post fracture. n= 8 mice. *: p<0.05, **: p<0.01, ***: p<0.001 vs *WT*.

### Bone defect healing does not require Prg4+ FAPs

Drill hole injury heals through a mechanism different from fracture. During its healing process, progenitors on the periosteal and endosteal surfaces directly differentiate into osteoblasts to fill the defect site. To explore the role of Prg4+ FAPs in this type of healing, we created a non-critical size defect in the right femurs of *Prg4ER/Td/Col1GFP* mice at 2 weeks post Tam injections. MicroCT analysis showed that the defect site is filled with new bone at 28 days (Fig. 7A, B). Since muscle was pushed aside to expose the periosteum during the injury, Prg4+ FAPs were also greatly expanded in nearby muscle on day 5 (Fig. 7Ci). However, they did not migrate to the cortical bone defect area (Fig. 7Cii). On day 14 when osteogenic cells were abundant in the newly formed woven bone at the defect area, Td+ cells remained mostly in the muscle (Fig. 7D). Some Td+GFP+ cells were located at the interface between muscle and the defect area (Fig. 7Di) and in the woven bone (Fig. 7Dii). At day 28, the newly formed cortical bone at the periosteum side contained Td+GFP+ cells, but they were rare (Fig. 7E). While these data demonstrated again that Prg4+ FAPs can become bone cells after injury, they also suggest that they do not contribute significantly to bone repair in the drill hole model.

**Figure 7.**
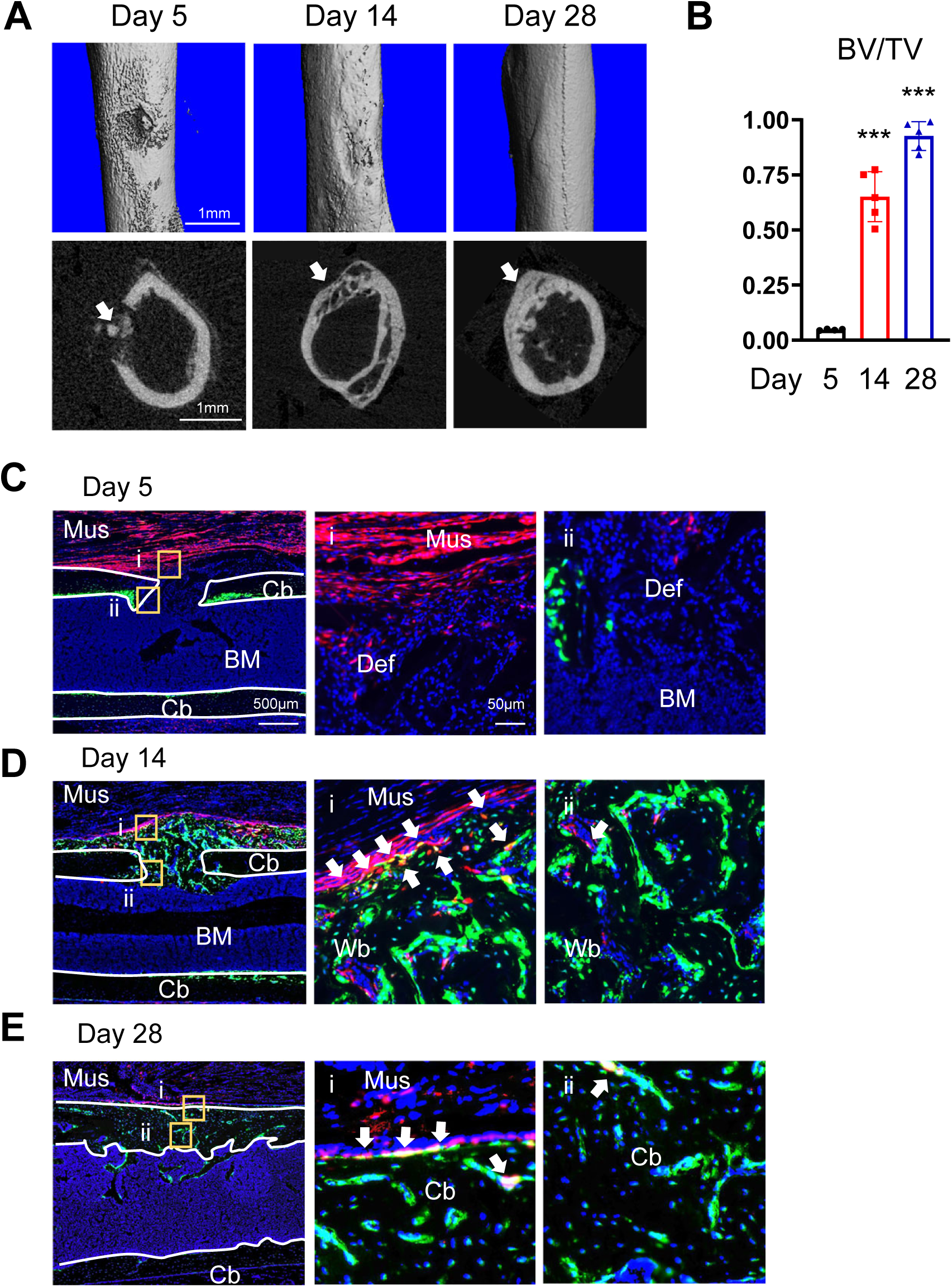
Prg4+ FAPs do not significantly contribute to the healing of cortical bone defect. (A) Representative sagittal (top) and transverse (bottom) cross-sectional microCT images of drill-hole defects in *Prg4ER/Td/Col1GFP* mice. Mice received Tam injections at 2 months of age followed by drill hole injury at right femurs 2 weeks later. Injured bones were harvested on days 5, 14, and 28 after injury for examination. Arrows point to the defect region. Scale bar: 1 mm. (B) Quantification of bone volume fraction (BV/TV) at cortical defect regions. n =5 mice/group. (C-E) Representative fluorescence image of *Prg4ER/Td/Col1GFP* femurs at days 5 (C), 14 (D), and 28 (E) post injury. The left panels are low magnification images (scale bar: 500 μm). Squares from those images are magnified at the right panel (scale bar: 50 μm). Arrows point to Td+GFP+ osteoblasts and osteocytes. Mus: muscle; Cb: cortical bone; Def: defect region; Wb: woven bone; BM: bone marrow. ***: p<0.001 vs *WT*.

## Discussion

In this study, we demonstrate that muscle FAPs are an active and indispensable contributor to bone regeneration by leveraging the ability of *Prg4-CreER* to specifically label a subset of muscle-resident FAPs, but not bone cells. When bone is injured, nearby skeletal muscle is also acutely damaged, a condition known to massively amplify FAPs. During repair, FAPs play a dual role. On the one hand, activated FAPs secrete cytokines and growth factors to promote MuSC proliferation and differentiation and to recruit immune cells for muscle repair (33). On the other hand, activated FAPs migrate to the hematoma and contribute to callus formation and bone regeneration (Fig. 8). After muscle injury, the number of FAPs peaks around 3-4 days (35), which coincides with hematoma formation and periosteum thickening post fracture (1). Our study revealed that FAP and their descendants are accumulated in the hematoma but not in the expanded periosteum at this time point. Shortly after, these cells differentiate into chondrocytes and osteoblasts in the outer part of the callus that is close to the muscle. In contrast, chondrocytes and osteoblasts in the inner part of the callus close to bone are exclusively derived from expanded periosteal mesenchymal progenitors. Although restricted by location, the contribution of FAPs to fracture callus is substantial because depletion of Prg4+ FAPs significantly reduces callus bone volume and delayed fracture healing. Eventually, after remodeling, FAPs evolve into periosteal mesenchymal progenitors and osteocytes at the repaired site. Those FAPs-derived bone cells now have the ability to respond to another injury in a similar way as other native periosteal mesenchymal progenitors. To our knowledge, this is the first report demonstrating that progenitors in a tissue can transform into progenitors in another tissue and differentiate into functional cells. Both FAPs and periosteal mesenchymal progenitors are organ-specific fibroblasts (36). While fibroblast plasticity is recognized within the same tissue, our finding reveals the plasticity of fibroblasts across different tissues.

**Figure 8.**
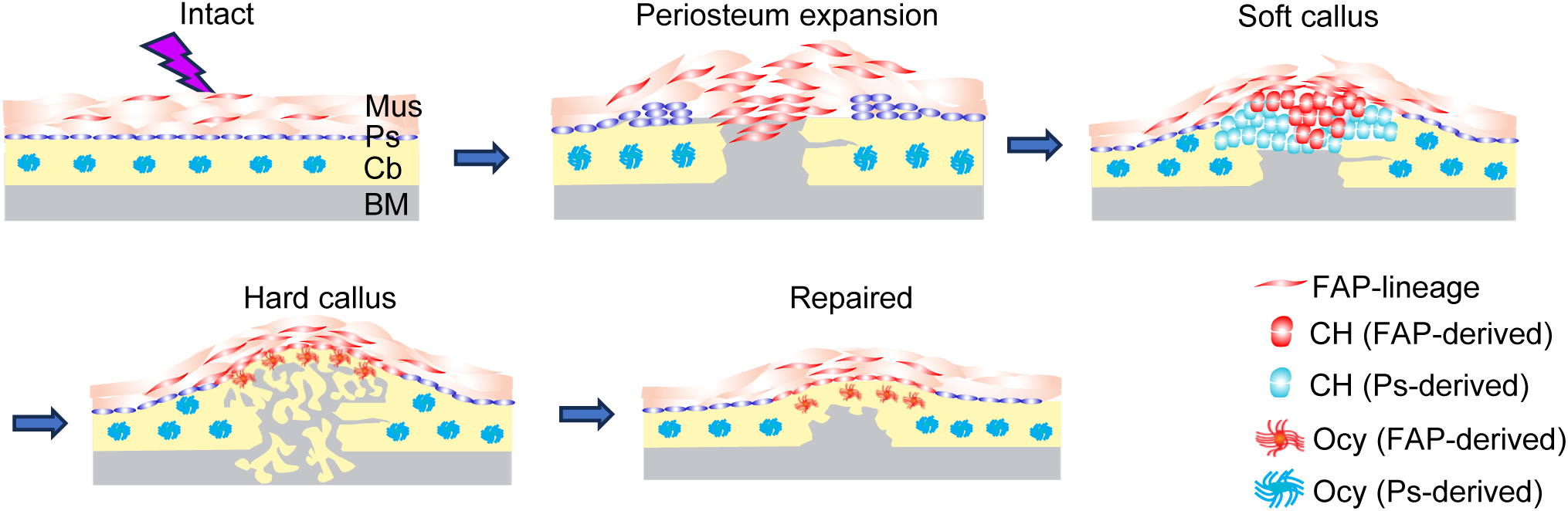
A schematic cartoon shows muscle FAP contribution to bone fracture healing. CH: chondrocyte; Ocy: osteocyte. Mus: muscle; Cb: cortical bone; Ps: periosteum; BM: bone marrow.

It is well known that muscle crosstalks with bone via mechanical loading as a contractile tissue attached to bone. The emerging theme from the past decade’s research is that both tissues act as secretory “paracrine” organs to regulate each other’s metabolism (37). Muscle secreted factors, so called “myokines” mainly derived from myoblasts, have negative or positive actions on bone. Meanwhile, bone-secreted factors, so-called “osteokines” mainly derived from osteoblasts and osteocytes, substantially influence muscle strength. Under injury scenarios, the communication from muscle to bone is more prominent than the communication in the opposite direction. Clinical observations and large animal studies have recognized the vascular role of muscle in bone regeneration for a long time (15, 38, 39). In particular, muscle flap coverage increases bone blood flow and the rate of osteotomy union compared to skin coverage, which has a poor vascular bed. Using muscle grafts, the Colnot group recently discovered that muscle FAPs, but not MuSCs, become a part of callus formed after fracture (18). Our study not only directly validates these findings but also uncovers a new crosstalk mechanism where bone injury transforms muscle FAPs into periosteal mesenchymal progenitors and osteocytes in the cortical bone. This unique type of cross-tissue trans-differentiation also occurs during tissue growth and homeostasis, and after muscle injury. However, the extent of contribution is much lower in those conditions compared to bone injury, particularly fracture.

As a whole, FAPs are traditionally characterized and isolated as Lin-Sca1+Cd34+ cells or Pdgfrα+ cells (35). We recently performed scRNA-seq analysis of intact tibial periosteum and callus at day 5 and day 10 from tibial fracture and discovered 6 clusters of mesenchymal subpopulations: mesenchymal progenitor cells (MPCs), proliferative progenitor cells (PPCs), early osteoblasts, osteoblasts, chondrocytes, and hypertrophic chondrocytes (40). Trajectory analysis revealed that MPCs are at the apex of mesenchymal lineage cells. Interestingly, MPCs are also labeled by Sca1, Cd34 and Pdgfra, the same markers defining FAPs. Mesenchymal progenitors are indistinguishable from fibroblasts (41).

Indeed, Sca1 and Cd34 are stem cell markers for fibroblast lineage cells across multi-organs and Pdgfra is a general marker for all fibroblast lineage cells (42). The similarity between periosteal MPCs and muscle-resident FAPs hints they might inter-convertible. Interestingly, we only observed that muscle-resident cells become bone-resident cells but did not find evidence supporting the opposite direction of plasticity.

Despite sharing the common fibroblast stem cell markers, a number of publications have revealed the heterogeneity of FAPs and suggested distinct roles of FAP subpopulations in supporting muscle development and pathological dystrophy (43). We found that Prg4+ FAPs overlap with Clu+ FAPs, a subpopulation identified in a recent study that displays high *in vitro* osteogenic differentiation ability (29). This feature is in line with our observation that Prg4+ FAPs become osteoblasts and osteocytes in the fracture callus. In addition to Clu+ FAPs, other 5 FAP subpopulations (Osr1+, Adam12+, Gap43+, Hsd11b1+, and Gli1+) were also identified in that scRNA-seq study. It would be interesting to explore whether multiple FAP subpopulations contribute to fracture healing and become periosteal mesenchymal progenitors. To do so, a key step is to construct FAP subpopulation-specific inducible *Cre* mouse lines for lineage tracing experiments.

The successful development of pharmaceutical therapies to promote bone healing has been fraught with challenges. Potential therapeutics that stimulate bone formation are limited to regulating Wnt and PTH/RANK pathways (romosozumab, teriparatide, and abaloparatide) and are indicated for the systemic building of bone mass rather than local bone regeneration or healing. The local stimulation of bone with BMP (as a therapeutic device) has demonstrated disappointing clinical results. In that context, research in this field has primarily focused on activating periosteal and endosteal musculoskeletal progenitor cells and has ignored cells in nearby muscle. Based on our study and others, we propose to expand future drug discovery to include those activating muscle-resident FAPs at the site of injury. Understanding signaling pathways that are injury specific (temporally and spatially), and that stimulate FAP proliferation and migration to the injury site upon fracture will shed new light on identifying novel drugs for non-unions and delayed unions, and perhaps for the acceleration of normal healing.

## Materials and Methods

### Analysis of scRNA-seq datasets

Pre-aligned scRNA-seq matrix files were acquired from GEO GSE164573 (mouse), GSE200234 (mouse), and GSE167186 (human). Standard Seurat pipeline (44) was used for filtering, normalization, variable gene selection, dimensionality reduction analysis, and clustering. Cell type was annotated according to the metadata from published datasets (18, 27).

### Animals study design

All animal work performed in this report was approved by the Institutional Animal Care and Use Committee (IACUC) at the University of Pennsylvania. In accordance with the standards for animal housing, mice were group housed at 23-25°C with a 12 hr light/dark cycle and allowed free access to water and standard laboratory chow.

*Prg4-CreER Rosa-tdTomato* (*Prg4ER/Td*) mice and *Scx-CreER Rosa-tdTomato* (*ScxER/Td*) were generated by breeding *Rosa-tdTomato* (Jackson Laboratory, 07909) mice (45) with *Prg4-CreER* mice (24) and *Scx-CreER* mice (Jackson Laboratory, 032132) (46), respectively. *Prg4-CreER Rosa-tdTomato Col1-GFP* (*Prg4ER/Td/Col1-GFP*) mice were generated by breeding *Prg4ER/Td* mice with *Col1a1*2.3-GFP* (Jackson Laboratory, 013134) mice (47). For cell ablation experiments, we bred *Prg4ER/Td* mice with *Rosa-DTA* mice (Jackson Laboratory, 009669) (48) to generate *Prg4ER/Td/DTA* mice. *Scx-GFP* mice (Jackson Laboratory, 032129) (49) were also used in our study. To induce Td expression or ablate Td+ cells, *Prg4ER/Td* or *Prg4ER/Td/DTA* mice received Tamoxifen (Tam) injections at 75 mg/kg for 5 days. Since we did not detect any gender difference in our experiments, both male and female mice at 8-12 weeks of age were used and comparison between different groups was performed using the same gender.

### Models of muscle and bone injury

To induce muscle injury, 10 μl Notexin (10 μg/mL, Latoxan; Accurate Chemical, TXL8104-100) was injected into mouse tibialis anterior (TA) muscle with an insulin syringe to induce acute muscle injury. To induce muscle injury with fatty degeneration, 50 μl 50% v/v glycerol (Sigma, G5516) in water was injected into mouse TA muscle. Both procedures were described in our previous publication (50). To perform fracture, mice received a closed transverse fracture on their right tibiae via a blunt guillotine with a pre-inserted intramedullary pin for stabilization as previously described (7). To perform drill-hole injury, we used a drill bit first followed by a 21-G needle to make a 0.8-mm diameter unicortical bone defect in the diaphysis part of mouse right femur as previously described (51).

### Micro-computed tomography (microCT) analysis

Tibiae harvested post fracture were scanned at the fracture sites by µCT 45 (Scanco Medical AG) at a 7.4-µm isotropic voxel size to acquire a total of 1098 µCT slices centering around the fracture site. A semi-automated contouring method was used to determine the callus perimeter and to analyze the callus outside the preexisting cortical bone. All images were first smoothed by a Gaussian filter (sigma = 1.2, support = 2.0) and then applied by a threshold corresponding to 30% of the maximum available range of image gray scale values to distinguish mineralized tissue from unmineralized and poorly mineralized tissue. Callus region surrounding cortical bone was contoured for trabecular bone analysis. Based on μCT images, day 56 fracture samples were assigned fracture healing scores according to an 8-point radiographic scoring system (52). This is a sum of scores from three categories: periosteal reaction (0–3), bone union (0–3), and remodeling (0–2). To analyze bone healing after drilling a hole, the contouring of the defect area was manually defined. A total of 150 slices were used to calculate bone volume fraction (BV/TV) in the cortical bone defect area.

### Mechanical testing

Fractured tibiae at day 56 post injury were tested immediately after isolation to prevent repeated freeze-thaw cycles and maintain structural integrity. They were carefully cleaned of soft tissue and potted into 5/16-inch round brass tubes (McMaster-Carr, Aurora, OH, USA) using low-viscosity veterinary bone cement (BioMedtrix, Whippany, NJ, USA). A standardized alignment guide, designed and 3D-printed specifically for this study, was employed to reproducibly expose the mid-shaft of the tibia and align the bone along the central axis. The potted tibiae were submerged in DPBS containing calcium and magnesium for 45 min to allow complete cement hardening and prevent evaporative losses. Subsequently, torsion testing was performed at a loading rate of 1°/second using a Mark-10 advanced torque testing system (Model TSTMH-DC) equipped with a 70 N·mm torque sensor (Mark-10, Copiague, NY, USA). Maximum torque (N·mm), torsional stiffness (N·mm/deg), and torsional energy to failure (N·mm·deg) were determined from the resulting torque-twist curves.

### Histology and immunohistochemistry

To examine bone and muscle together, tibiae along with surrounding muscle were fixed in 4% paraformaldehyde (PFA) for 24 hr, decalcified in 10% Ethylenediaminetetraacetic acid (EDTA) for 4 weeks, dehydrated in 30% sucrose, embedded in optimal cutting temperature (OCT) compound, and sectioned longitudinally at 6 μm in thickness using cryofilm tape (Section Lab, Hiroshima, Japan). For immunostaining, sections were incubated with rabbit anti-Osterix (Abcam, ab22552), rat anti-Cd45 (Biolegend, 103101), or rabbit anti-Collagen II (abcam, ab34712) at 4°C overnight followed by Alexa Fluor 488 anti-rat (Invitrogen, A21208) or Alexa Fluor 647 anti-rabbit (Abcam, ab150075) secondary antibody incubation for 1 hr at RT. For osteocyte staining, sections were stained with DyLight™ 488 Phalloidin (Cell signaling, 12935). For H&E imaging, sections adjacent to those for fluorescence imaging were stained by Harris Hematoxylin solution and 0.5% Eosin Y and mounted in 50% glycerol and scanned immediately.

To examine muscle tissue only, TA muscles were fixed in 4% PFA, dehydrated in 30% sucrose, embedded in OCT compound, and sectioned cross-sectionally at 10 μm in thickness. For immunofluorescence staining, sections were blocked by 3% bovine serum albumin (BSA) for 30 min and incubated with rat anti-Sca1 (BioLegend, 122501), mouse anti-Pdgfrα (Santa Cruz, SC-398206), or rabbit anti-Perilipin (Cell Signaling, 9349) antibodies at 4°C overnight, followed by secondary antibody incubation for 1 h. To label the extracellular matrix, sections were stained with CF®640R Wheat germ agglutinin (WGA, Biotium, 29026-1) for 1 h.

For imaging the morphology of muscle post muscle injury, TA muscle was fixed in 4% PFA and processed for paraffin sections and H&E staining.

For RNA FISH experiment, we adopted in situ hybridization chain reaction (HCR) approach (Molecular Instruments, Los Angeles, CA). Briefly, cryosections were processed and stained by probes against *Prg4* (NM_001110146), mRNAs according to the manufacturer’s protocol (HCR™ RNA-FISH protocol for fresh frozen or fixed frozen tissue sections).

All sections were scanned by Axioscan (Carl Zeiss MicroImaging, Göttingen, Germany).

We used ImageJ with standardized thresholding and cell-counting protocols to quantify Prg4-lineage cells in the TA muscle and tibial periosteum. From each mouse sample, one section with the largest muscle area was selected for analysis. TA muscle area and the length of periosteum next to the TA muscle were measured. Td+ cells within the TA muscle and in the periosteum were counted to calculate Td+ cells per muscle area and Td+ cells per periosteum length.

### CFU-F assay

To isolate FAPs for CFU-F assay, hindlimb muscles (quadriceps, gastrocnemius, and tibialis anterior) were dissected, trimmed off surrounding ligament tissues, minced with scissors, and enzymatically dissociated with 0.1% collagenase Type II (Gibco, 17101015) in DMEM on shaker for 2 hr in 37°C incubator. Digested cells were filtered through a 40 μm cell strainer and seeded at 3×10^4^ cells per T25 flask in growth medium (α-MEM supplemented with 15% FBS, 0.1% β-mercaptoethanol, 20 mM glutamine, 100 IU/ml penicillin, and 100 µg/ml streptomycin) for 7 days before counting CFU-F number under the fluorescence inverted microscope (Leica, Germany) using bright field and fluorescence channel.

Periosteum cells were harvested as described previously (7). Intact tibiae or tibiae harvested at day 56 post fracture were dissected free of surrounding tissues and both epiphyseal ends were sealed with 3% agarose gel. The remaining bone fragments were digested in 2 mg/ml collagenase A and 2.5 mg/ml trypsin. Cells from the first 3 min of digestion were discarded and cells from a subsequent 30 min of digestion were collected and seeded in the growth medium at 1x10^6^ cells per T25 flask for 7 days before counting CFU-F number under the fluorescence inverted microscope (Leica, Germany) using bright field and fluorescence channel.

### FACS analysis

To isolate callus cells, after trimming surrounding muscles, the callus was cut by a scalpel, minced with scissors, and enzymatically dissociated with 2 mg/mL collagenase A (Roche Diagnostics) and 2.5 mg/mL trypsin dissolved in Dulbecco’s PBS for 2 hr, followed by filtering through a 40 μm cell strainer. Cells from muscle or callus were stained with fluorescence labeled primary antibodies, including FITC anti-Cd45 (Biolegend, 147709), FITC anti-Cd31 (Biolegend, 160211), FITC anti-Ter119 (Biolegend, 116205), Brilliant Violet 605 anti-Sca1 (Biolegend, 108113), BV421 anti-Cd140a (Pdgfrα) (Biolegend, 135923) for 1 hr at 4°C, washed in flow buffer (5% BSA, 2mM EDTA in HBSS) and filtered through a 35 μm cell strainer tube (Falcon). Cells were then analyzed using a BD LSRFortessa flow cytometer.

### Statistical analyses

Data are expressed as means ± standard deviation and analyzed by t-tests or one-way analysis of variance with Tukey post-test for multiple comparisons using Prism software (GraphPad Software, San Diego, CA). For cell culture experiments, observations were repeated independently at least three times with a similar conclusion, and only data from a representative experiment are presented. Values of p < 0.05 were considered significant.

## Supporting information

Supporting text Figures S1 to S11 Legends for Figures S1 to S11

## Acknowledgments

We thank Dr. Maurizio Pacifici at Children’s Hospital of Philadelphia and Dr. David Rowe at the University of Connecticut for providing us *Prg4-CreER* mice. We also thank Dr. Alice Huang at Columbia University for advice on studying tendon cell contribution to fracture healing. MicroCT Imaging Core at Penn Center for Musculoskeletal Disorders (PCMD) for their assistance with microCT analysis. This study was supported by NIH grants NIH/NIA R01AG069401 (to L.Q.) and P30AR069619 (to Penn Center for Musculoskeletal Disorders).

