## Supporting text Figures S1 to S11 Legends for Figures S1 to S11 for "Prg4+ fibro-adipogenic progenitors in muscle are crucial for bone fracture repair"

**This PDF file includes:**

Supporting text
Figures S1 to S11
Legends for Figures S1 to S11

**Figure S1**

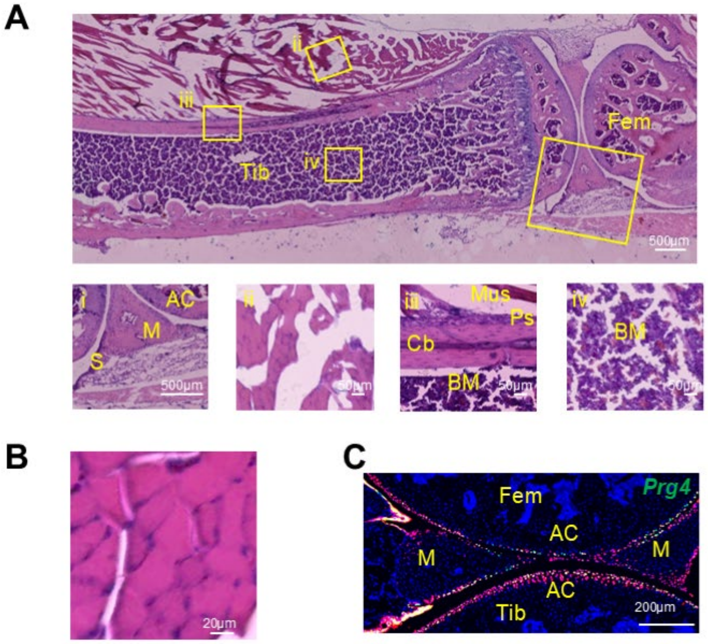


**Figure S1.** Prg4 expression labels a subset of FAPs in skeletal muscle.

(A) H&E staining of a section adjacent to the one used for fluorescence image in Fig. 1A. Top panel: a low magnification image containing tibia (Tib), femur (Fem), and joint tissue. Scale bar: 500 µm. Bottom panel, squares in the top panel are magnified to show joint (i, scale bar: 500 µm), muscle (Mus, ii, scale bar: 50 µm), cortical bone (Cb, iii, scale bar: 50 µm), and bone marrow (BM, iv, scale bar: 50 µm). M: meniscus; S: synovium; AC: articular cartilage; Ps: periosteum.

(B) H&E staining of a section adjacent to the one used for fluorescence image in Fig. 1B. Scale bar: 20 µm.

(C) Fluorescent image of a knee joint stained for *Prg4* mRNA by RNA FISH. Scale bar: 200 µm.

### Figure S2

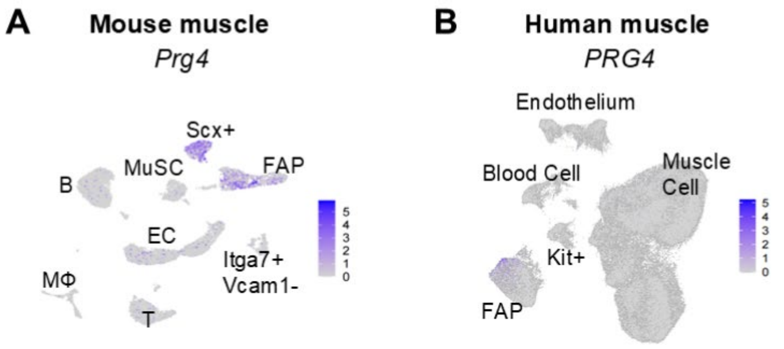


**Figure S2.** The expression level of *Prg4* in muscle single-cell dataset.

(A) The UMAP plot of *Prg4* expression in mouse muscle cells. B: B cell; MΦ: macrophage; EC: endothelium cell; MuSC; muscle stem cell.

(B) The UMAP plot of *Prg4* expression in human muscle cells.

**Figure S3**

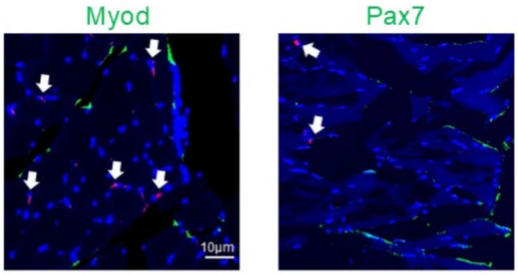


**Figure S3.** Prg4+ FAPs are not MuSCs.

*Prg4ER/Td* mice at 2 months of age received Tam injections on days 1-5, and their TA muscle was harvested on day 14 for crossectional cryosections, followed by Myod or Pax7 immunostaining. Arrows point to Td+ FAPs. Scale bar: 10 µm.

**Figure S4**

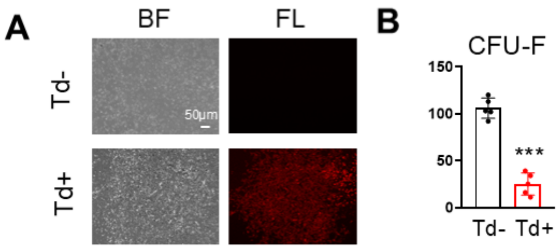


**Figure S4.** Prg4+ cells in muscle proliferate in culture.

(A) CFU-F assay of muscle cells from *Prg4ER/Td* mice shows a Td- colony (top) and a Td+ colony. BF: brightfield; FL: fluorescent light. Scale bar: 50 μm.

(B) Td+ and Td- CFU-F colonies were counted from 3 million digested muscle cells.

***: $p<0.001$, n= 5 mice.

**Figure S5**

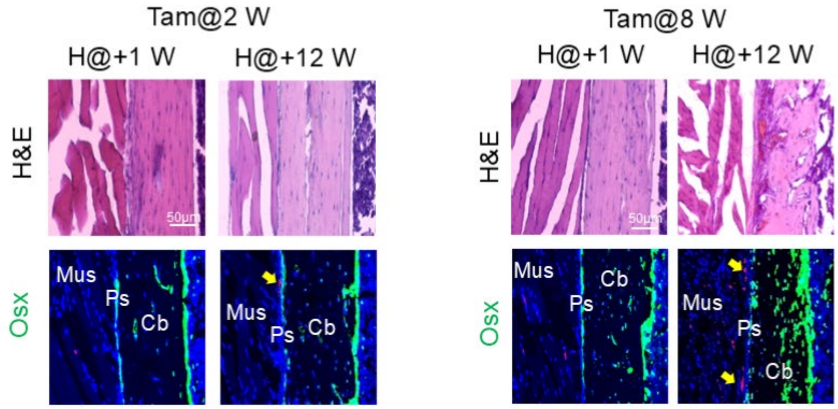


**Figure S5.** Prg4+ FAPs give rise to periosteal cells during muscle growth and maintenance. Mice at 2 or 8 weeks of age received Tam injections. Their tibiae and surrounding muscle were harvested at 1 and 12 weeks later for H&E images and fluorescence images with Osterix straining of adjacent *Prg4ER/Td* tibial sections. Arrows point to Td+ cells in the fibrous layer. Scale bar: 50 μm.

### Figure S6

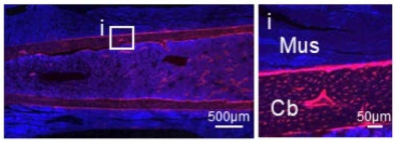


**Figure S6.** Col2-lineage bone cells do not become muscle cells.

Represent fluorescence image of tibiae and surrounding muscle from *Col2/Td* mice harvested at 2 months of age. A square in the low mag image at the left (Scale bar: 500 μm) is magnified at the right (**I**, scale bar: 50 μm).

**Figure S7**

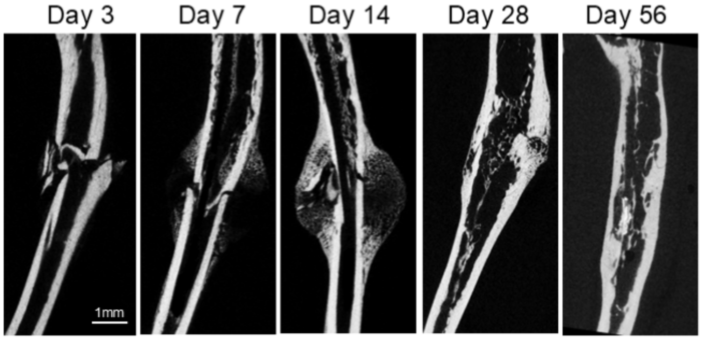


**Figure S7.** Representative microCT images of tibiae at different time points post fracture surgery.

Scale bar: 1 mm.

### Figure S8

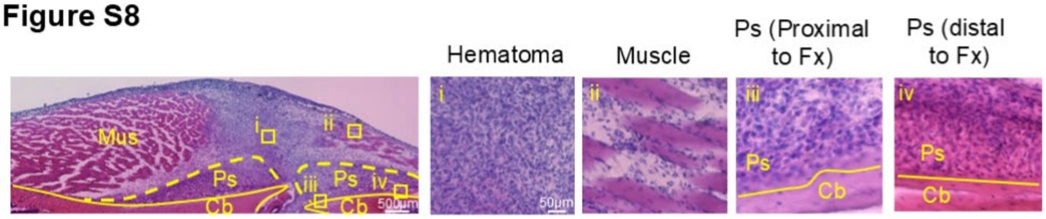


**Figure S8.** H&E staining of a section adjacent to the one used for fluorescence images in Fig. 4B.

**Figure S9**

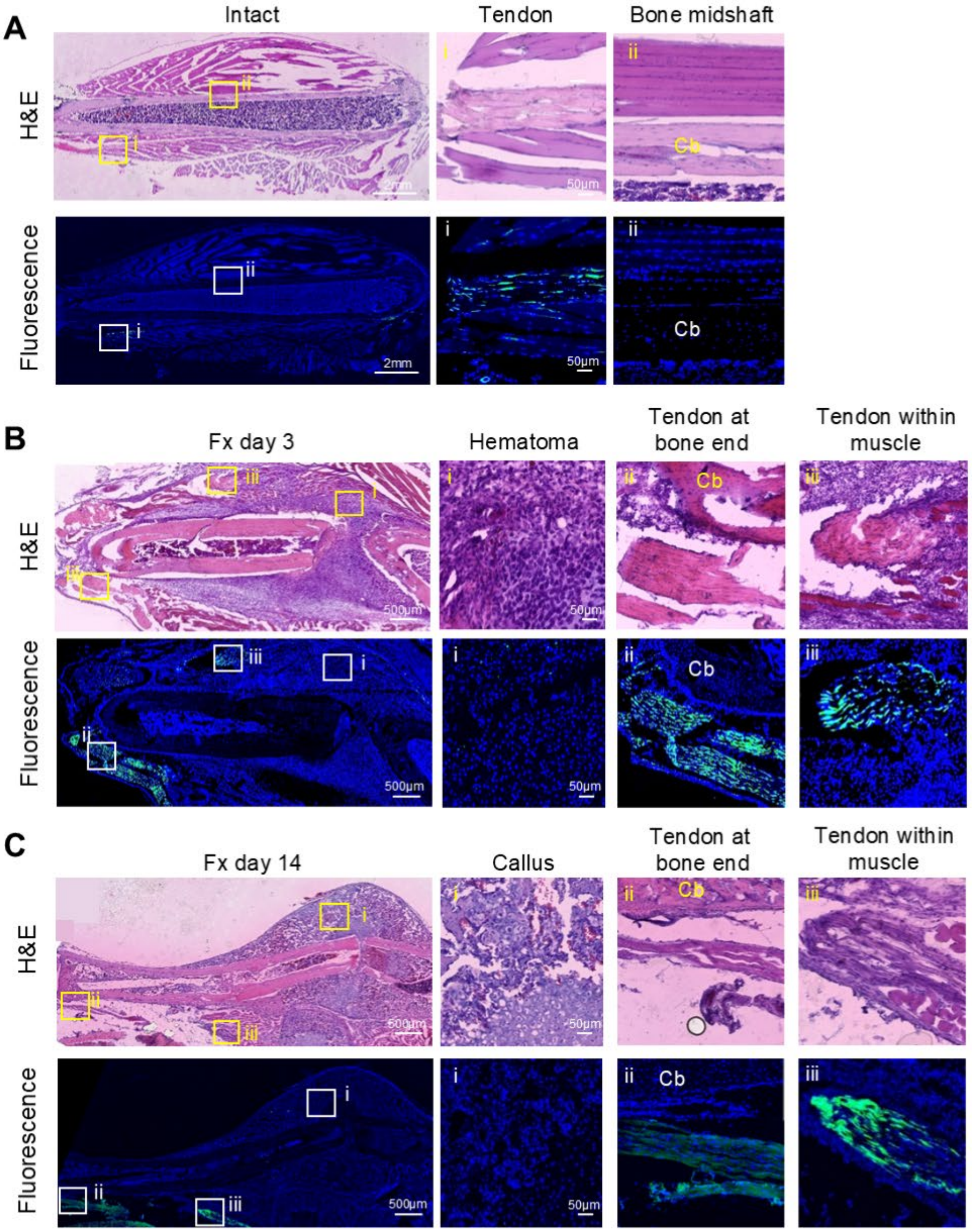

A
Intact
Tendon
Bone midshaft
H&E
Fluorescence
i
ii
Cb
2mm
50µm
B
Fx day 3
Hematoma
Tendon at bone end
Tendon within muscle
iii
500µm
C
Fx day 14
Callus

**Figure S9.** Tendon remains relatively intact after fracture.

(A) H&E and fluorescence images of adjacent tibia and muscle sections from 2-month-old *Scx-GFP* mice. Left panel: a low magnification tibiae image. Scale bar: 2 mm. Right panel, squares in the left panel are magnified to show tendon (i, scale bar: 50 μm), bone midshaft (ii, scale bar: 50 μm). Cb: cortical bone.

(B-C) H&E and fluorescence images of adjacent tibia and muscle sections from 2-month-old *Scx-GFP* mice at days 3 (B) and 14 (C) after fracture. Left panel: low magnification images of the fracture site. Scale bar: 500 μm. Right panel, squares in the left panel are magnified to show callus (i, scale bar: 50 μm), tendon at bone ends (ii, scale bar: 50 μm), and tendon within muscle (iii, scale bar: 50 μm).

**Figure S10**

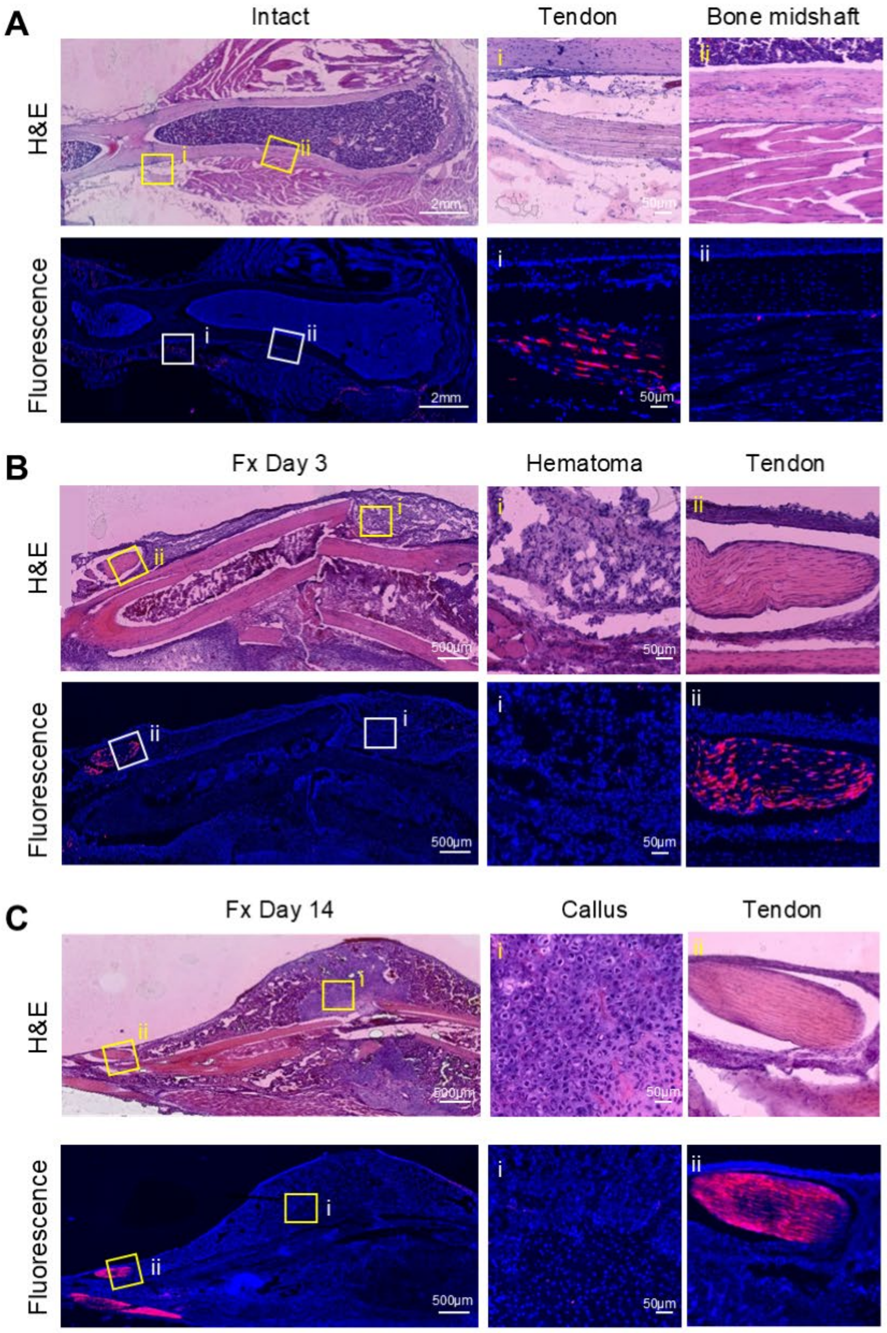
A
Intact
Tendon
Bone midshaft
H&E
Fluorescence
i
ii
2mm
50μm
B
Fx Day 3
Hematoma
Tendon
500μm
C
Fx Day 14
Callus

**Figure S10.** Tendon cells do not contribute to callus formation.

(A) H&E and fluorescence images of adjacent tibia and muscle sections from *ScxER/Td* mice. Two-month-old mice received Tam injections on days 1-5, followed by tissue harvest on day 14. Left panel: low magnification tibia images. Scale bar: 2 mm. Right panel, squares in the left panel are magnified to show tendon (i, scale bar: 50 μm), bone midshaft (ii, scale bar: 50 μm). Cb: cortical bone.

(B-C) H&E and fluorescence images of adjacent tibia and muscle sections from *ScxER/Td* mice at days 3 (B) and 14 (C) after fracture. Mice received fracture at 2 weeks post Tam injections. Left panel: low magnification images of the fracture site. Scale bar: 500 μm. Right panel, squares in the left panel are magnified to show callus (i, scale bar: 50 μm), tendon at bone end (ii, scale bar: 50 μm), and tendon within muscle (iii, scale bar: 50 μm).

### Figure S11

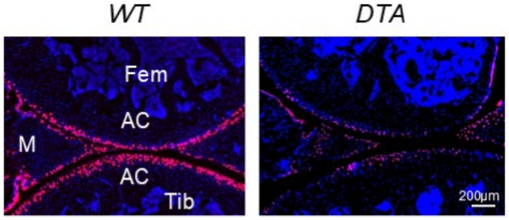


**Figure S11.** Ablation of Prg4+ cells is successful in the knee joint.

Representative fluorescence image of *Prg4ER/Td* (*WT*) and *Prg4ER/Td/DTA* (*DTA*) knee joints.

Fem: femur; Tib: tibia; M: meniscus; AC: articular cartilage. Scale bar: 200 μm.
